# MTX-COVAB, a human-derived antibody with potent neutralizing activity against SARS-CoV-2 infection *in vitro* and in a hamster model of COVID-19

**DOI:** 10.1101/2020.12.01.406934

**Authors:** Simone Schmitt, Marcel Weber, Matthias Hillenbrand, Jemima Seidenberg, Andreas Zingg, Catherine Townsend, Barbara Eicher, Justina Rutkauskaite, Peggy Riese, Carlos A. Guzman, Karsten Fischer, Christoph Esslinger

## Abstract

Fast track microfluidic screening of the antibody repertoires of 12 convalescent COVID-19 donors comprising 2.8mio antibodies yielded MTX-COVAB, a human-derived monoclonal antibody with low picomolar neutralization IC50 of SARS-CoV-2. COVAB neutralization potency is on par with the Regeneron cocktail as demonstrated in a comparative neutralization assay. MTX-COVAB shows strong efficacy *in vivo* and binds to all currently identified clinically relevant variants of SARS-CoV-2. MTX-COVAB completes GMP manufacturing by the end of this year and will be tested in the clinic in March 2021.

## Introduction

The development of recombinant antibody therapeutics for the prevention and treatment of viral infections from convalescent donors constitutes a straightforward counter measure in the current SARS-CoV-2 pandemic. We embarked on such an approach in March 2020 recruiting donors with mild to moderate and moderate to severe symptoms locally from the Zurich metropolitan region.

Screening for potentially virus-neutralizing antibodies was performed using our proprietary DROPZYLLA® technology, a droplet microfluidic single-cell-based antibody discovery platform. In brief, we copied the antibody genetic information of single memory B cells from each of the 12 donors into 293 HEK cells while maintaining heavy and light chain pairing of the original B cells. The antibodies were expressed in the HEK cells as membrane-bound full-length IgG (mIgG-HEK) to enable antigen-specific sorting. Our initial screening strategy consisted of isolating binders to the receptor-binding domain (RBD) of the SARS-CoV-2 spike protein using fluorescence labeled RBD followed by neutralization assays. Due to a high ratio of non-neutralizing binders, we then switched to screening for those mIgG-HEK that bound to RBD but blocked the binding of ACE2. For this we used ACE2 labeled with a second fluorochrome. Most antibodies that were identified from this screening proved to also neutralize SARS-CoV-2 infection in later assays. The time from blood donation to the identification of antibodies of interest was between 3 to 6 weeks.

As a secondary screening, binders were screened for neutralizing capacity in a lentivirus-based pseudovirus assay and antibodies with picomolar neutralizing activity were subsequently tested in plaque assays using a clinical isolate of SARS-CoV-2. Additional tests for binding and neutralization of variants of SARS-CoV-2 and developability (stability and yield) were then performed to identify our development candidate MTX-COVAB.

## Results & Discussion

### MTX-COVAB, a strong neutralizer of SARS-CoV-2 infection *in vitro*

As a major determinant of its anti-infective activity, the neutralization capabilities of MTX-COVAB towards a clinical isolate of SARS-CoV-2 was assessed in vitro. After performing a ranking of the candidate antibodies using the pseudovirus system, we submitted the 4 best antibodies to a neutralization plaque assay with replication competent wild type virus. We observed a linear relationship to the results obtained with the Pseudovirus test. The best performing antibody in both assays, CoVAb36 was selected as lead antibody since it combined excellent virus neutralization activity (see below figure 1) with favorable developability attributes (data not shown).

**Figure 1.**
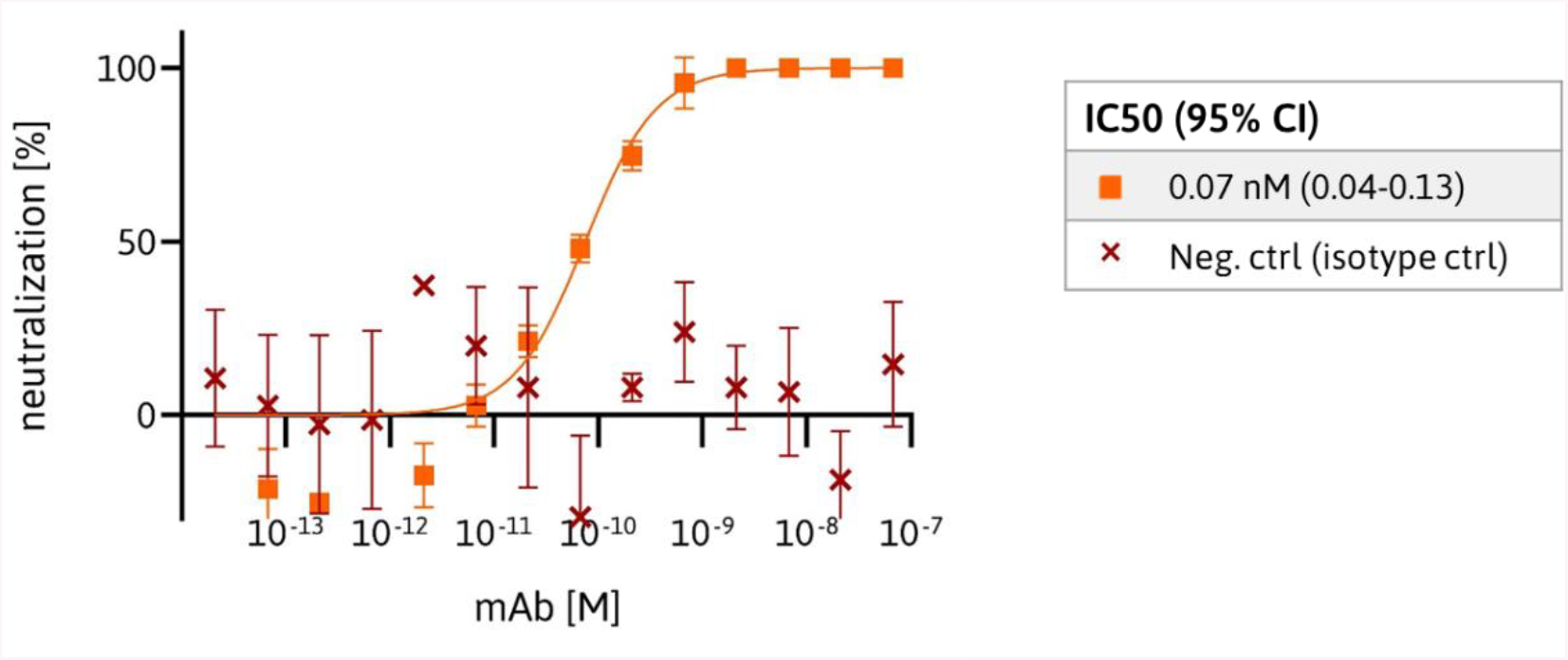
Neutralization of wild type SARS-CoV-2 by MTX-COVAB. The anti-SARS-CoV-2 antibody and a control antibody were tested for their capability to neutralize the SARS-CoV-2 virus by conducting plaque assays. Data are depicted as percentage of neutralization relative to the average number of plaques in the negative control (0%) and no plaques visible (100%). Data were analyzed and IC50 values were determined using the [Inhibitor] vs. response -- Variable slope” fitting model of GraphPad Prism 8. Symbols shown are mean of triplicates and error bars are SD.

**Figure 2:**
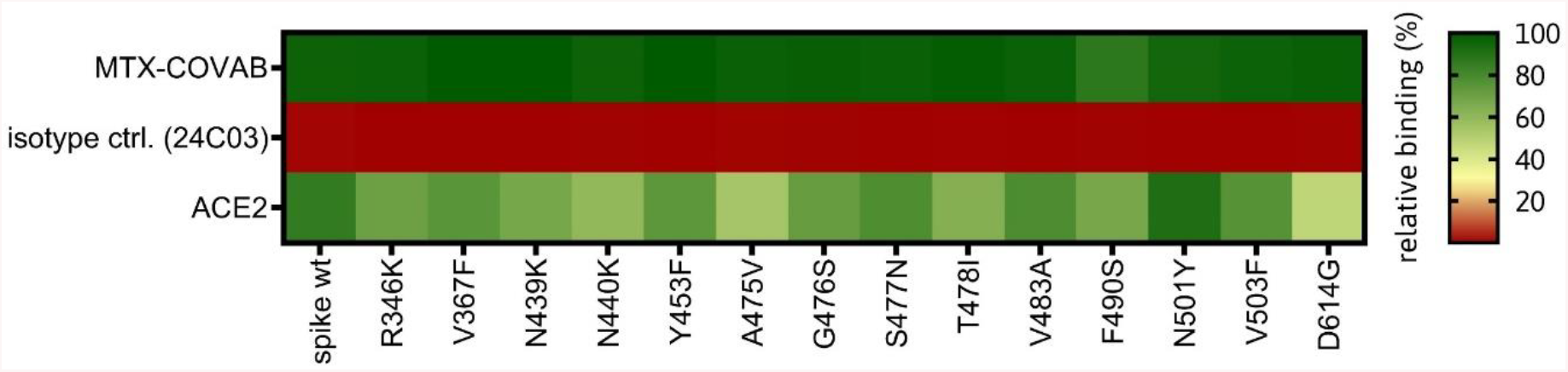
Binding of MTX-COVAB to relevant SARS-CoV-2 spike mutations. Mutants of SARS-CoV-2 Spike protein were expressed on HEK293T cells and cells were incubated with MTX-COVAB, an irrelevant control antibody (24C3) and the cellular receptor of SARS-CoV-2, ACE2. Relative binding strength is indicated by colors where dark green represents strong binding and dark red no binding.

### MTX-COVAB binds to naturally occurring variants of SARS-CoV-2

We generated 293 HEK cells expressing selected, naturally occurring variants of the SARS-CoV-2 spike protein to assess the binding of MTX-COVAB in comparison to the wildtype SARS-CoV-2 Spike protein. All of the mutants contain a single mutation compared to the original Wuhan (wt) strain.

MTX-COVAB bound to all of the variants in a similar fashion. In contrast, ACE2, the natural receptor of the SAR-CoV-2 spike showed a generally much lower binding strength which was expected due to its significantly lower affinity to the spike compared to MTX-COVAB. The more heterogenic pattern may also inform on the relevance of a given variant since attachment to ACE2 is a mandatory prerequisite for a successful infection.

### MTX-COVAB shows strong *in vivo* activity in the Syrian hamster model

The Syrian golden hamster model for SARS-CoV-2 infection was chosen due to the increased relevance of this model to the human disease in terms of strong lung pathology and a dramatic loss in body weight, which was in turn our main clinical read out for effectivity of the antibody (Imai et al., Baum et al. Kreye et al.). We assessed MTX-COVAB both as a prophylactic and a treatment as outlined in Figure 3. When used as a prophylaxis, MTX-COVAB prevented weight loss even at day 1 after virus inoculation. With all doses tested, including the lowest dose administered (1 mg/kg), a gain of weight was visible in a dose-dependent manner that reached a plateau at 10 mg/kg body weight. When used as a treatment, a dose-dependent recovery from weight loss could be observed. This therapeutic effect was most striking at day 7, where a maximal weight loss of 15% was observed with Placebo compared to less than 3% with the two higher doses of 25 and 50 mg/kg. In summary, we could demonstrate that MTX-COVAB is able to reduce the clinical symptoms of infection both in a therapeutic and prophylactic setting.

**Figure 3.**
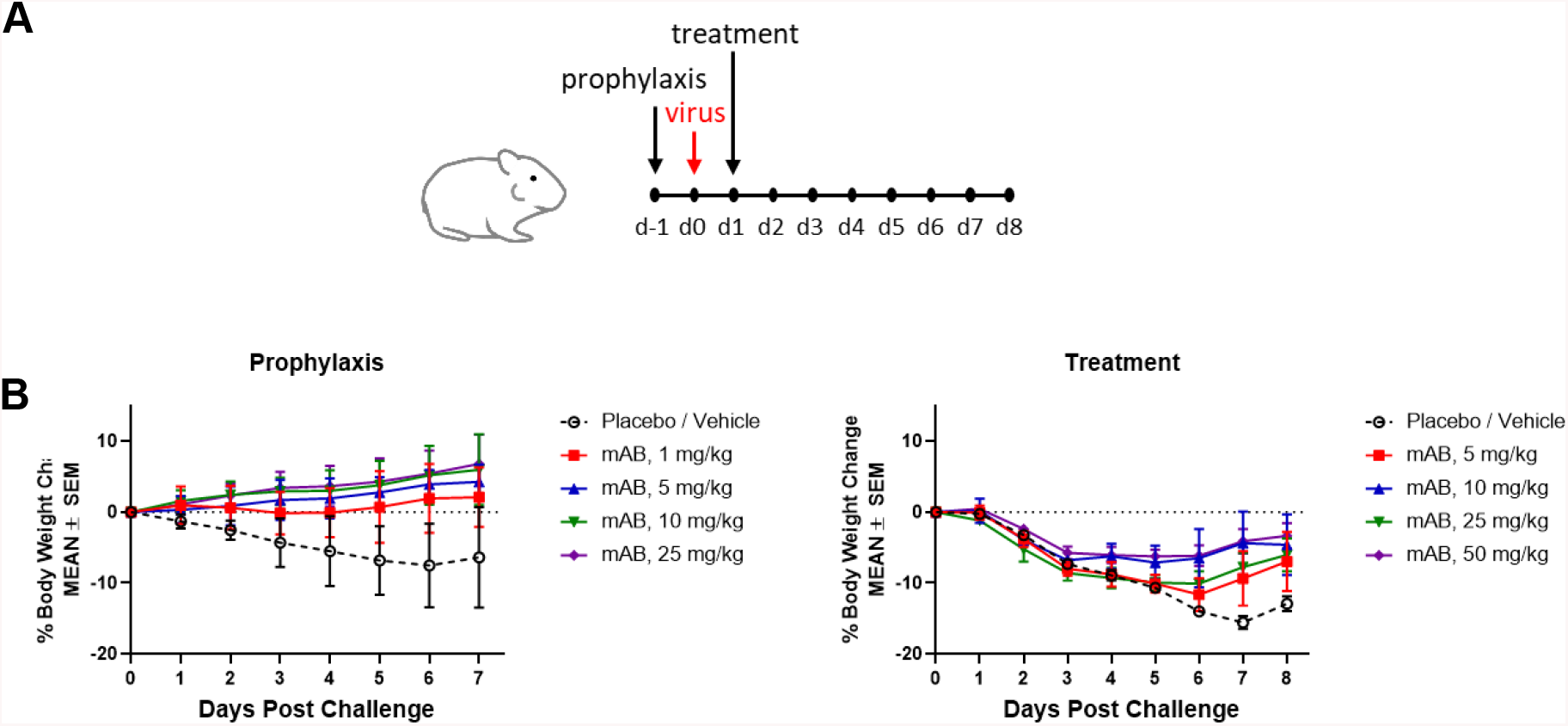
*In vivo* efficacy of MTX-COVAB for prophylaxis and treatment of SARS-CoV-2 infection. **(A)**Outline of study in the Syrian hamster model **(B)** Clinical readout: effect of COVAB on weight loss in prophylaxis and treatment.

### ADCC, ADCP & CDC may be additional mechanisms of action of MTX-COVAB

Fc-mediated effector functions such as antibody dependent cellular cytotoxicity, antibody dependent cellular phagocytosis and complement dependent cytotoxicity (ADCC, ADCP and CDC) could be important components of the anti-infectious activity of SARS-CoV-2-neutralizing antibodies.

For this reason, stable SARS-CoV-2-spike protein transfected 293 HEK cells were generated as targets. In order to assay MTX-COVAB for ADCC, purified human NK cells were used as effector cells. In this assay, MTX-COVAB showed a strong, dose-dependent, ADCC effect (Fig. 4 A). Using a Fc-γRIIIa-dependent reporter cell assay a similar result was obtained, indicating that ADCC was mediated via this Fc-receptor (data not shown).

**Fig 4.**
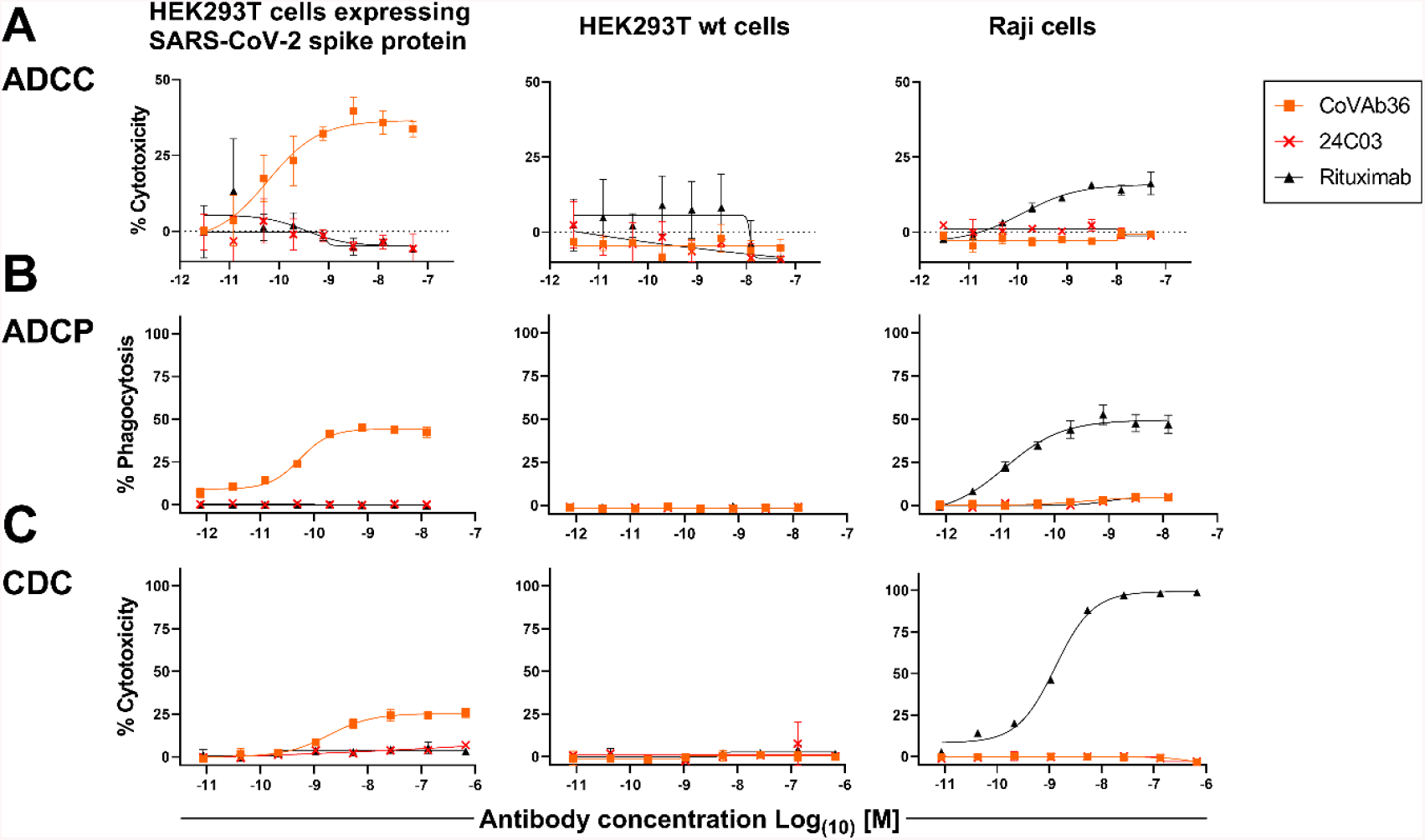
Anti-SARS-CoV2-Spike antibody MTX-COVAB induces specific ADCC, ADCP and CDC. HEK293T cells expressing SARS-CoV-2 Spike protein, parental HEK293T cells or Raji (CD20^+^) cells were mixed with (A) NK cells for ADCC, (B) with PBMCs for ADCP and (C) with 20% human serum for CDC. A titration of anti-SARS-CoV2-Spike protein antibody MTX-COVAB, the IgG1 isotype control 24C03 or anti-CD20 antibody Rituximab was added to cells and incubated overnight. Cytotoxicity on target cells (A &C) was assessed using SYTOX™ Blue Dead Cell Stain, whereas phagocytosis (B) was assessed by the uptake of target cells by CD14+ monocytes. The baseline was defined as the average percent cytotoxicity/phagocytosis of the target/effector cell co-culture in absence of antibody. Each condition was tested in triplicate. Like Rituximab, which only affected Raji cells, CoVAb36 only showed ADCC, ADCP and CDC on SARS-CoV-2 expressing target cells, but not on untransfected cells.

When testing for ADCP using whole human PBMC as effector cells and measuring the uptake of target cells into cells of the monocyte lineage, MTX-COVAB again showed a dose-dependent ADCP (Fig. 4 B).

CDC by MTX-COVAB was less pronounced but still measurable (Fig. 4C).

Taken together, MTX-COVAB mediates strong Fc-receptor-dependent cytotoxicity, which may well constitute an additional mechanism of action against the virus. A caveat for this is the fact that the target cells we used were stably transfected cells that may or may not mimic the *in vivo* situation in an infected individual. The only experimental evidence for ADCC against corona virus infected cells that we could identify is described in Holmes et al. Br. J. exp. Path. I986.

### Comparison to benchmark antibodies currently in the clinic

In an attempt to benchmark the virus neutralizing activity of MTX-COVAB to one of its peers, we compared it side-by-side with the two antibodies that form Regeneron’s cocktail (Hansen et al. Science 2020) using our pseudovirus assay. As shown in Figure 5, MTX-COVAB shows a neutralization capacity that is on par with that obtained with the better of the two antibodies as well as with the cocktail.

**Figure 5.**
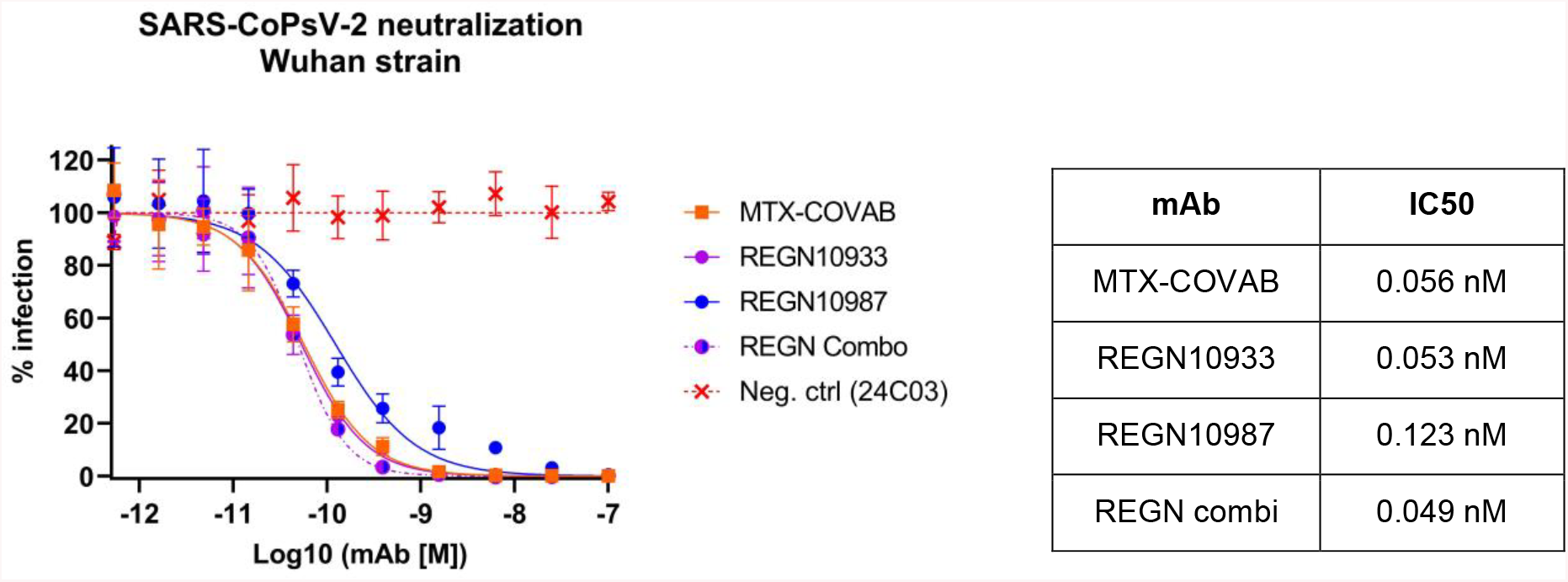
Comparison MTX-COVAB with REGN-CoV-2 antibodies and cocktail in SARS-CoV-2-Pseudovirus (SARS-CoPsV-2) neutralization assay. Indicated are percent infection of HEK-ACE2 cells. While 100% reflects the average signal from isotype control antibody (negative control, mAb 24C03) and 0% reflects the background signal. Table indicates half maximal inhibitory concentration (IC50) for each of the antibodies and for the REGN combi (REGN10933 and REGN10987 in a mixture of equal concentrations) IC50 values were determined using the [Inhibitor] vs. normalized response -- Variable slope” fitting model of GraphPad Prism 8. Symbols shown are mean of triplicates and error bars show SD.

## Conclusion

MTX-COVAB potently neutralizes SARS-CoV-2 in vitro, retains binding to naturally occurring variants of the SARS-CoV-2 spike and also shows potent *in vivo* activity against SARS-CoV-2 infection in the Syrian hamster model. MTX-COVAB is an unmodified human IgG1 antibody and retains the ability to mediate ADCC, ADCP and CDC.

## Methods

### Donors & Ethics

Donors were recruited in accordance with the laws of Switzerland and under ethics approval BASEC-2016-01260 of the Cantonal Ethics Commission of Zurich.

### Antibody discovery

Memory B cells from peripheral blood were used to prepare antibody repertoire expression libraries by cloning the immunoglobulin light chain and heavy chain variable region into an expression cassette providing the human immunoglobulin constant heavy region combined with a transmembrane domain derived from human CD8 to allow for mammalian cell display of the antibodies. Screening of the antibody libraries was performed after transduction of the library in HEK 293T cells by antigen-specific sorting using fluorescently labelled SARS-CoV-2 Spike RBD. This sort yielded RBD-specific-antibody-expressing HEK cell clones. Clone specificity was confirmed in analytical FACS, testing for high-affinity binding of cell-membrane expressed antibodies to purified SARS-CoV-2 Spike RBD protein and simultaneously absence of RBD-binding to purified huACE2. Antibody clones positive for RBD binding and negative for huACE2 binding were sequenced and sub-cloned into expression vectors for soluble IgG expression and expressed after transient transfection in HEK 293F. Antibodies were then purified via Protein A for further characterization in the various assays.

### Preparation of pseudo-typed SARS-CoV-2 (SARS-CoPsV-2)

Lentivirus-based pseudo-typed virus particles with SARS-CoV-2 Spike protein or a Spike protein mutant, replacing the VSV-G gene, were cloned and produced at MTX. As a reporter of infection, a transfer vector encoding a constitutively expressed, secreted luciferase was co-transfected during production of pseudo-typed virus batches.

### Neutralization of SARS-CoPsV-2

HEK293T cells stably expressing full length huACE2 (aa1-805) were seeded and left to adhere. Serial antibody dilutions were prepared and added to the adherent cells. Directly thereafter, SARS-CoPsV-2 carrying a secreted luciferase reporter gene was added and the mix was incubated for 3 days. After this period, the amount of secreted luciferase in the supernatant was determined by luciferase assay (NanoGlo® Luciferase Assay, Promega, #N1130 for the qualitative analysis of infected cells. Data were analysed and IC50 values were determined using the “log [Inhibitor] vs. normalized response -- Variable slope” fitting model of GraphPad Prism 8.

### Binding to variants of SARS-CoV-2 spike

Variants of the SARS-CoV-2 spike protein were generated by gene synthesis and expressed in HEK293T cells. Cells were incubated with MTX-COVAB or an isotype control antibody (24C03). Binding of antibodies was detected using a fluorescence labeled anti human IgG. Binding to ACE2 was measured by incubating cells expressing the spike protein variants with recombinant ACE2 containing a His Tag. Bound ACE2 was then detected using a fluorescence labeled anti His Tag antibody. The fluorescence stained cells were analyzed by flow cytometry.

### Neutralization of wild type SARS-CoV-2

At day 0, Vero E6 cells were seeded with a concentration of 1*105 cells/well in 300 μl medium. At day 1, the antibodies were serially diluted using a 1:3.16 dilution starting with 1 or 10 μg/ml in a total volume of 300 μl/well. Subsequently, the antibodies were mixed with SARS-CoV-2 virus (1800 PFU/ml) and incubated for 1h at 37°C. Then, media from the Vero E6 cells was removed and the cells were incubated with the antibody-virus mix for 1h. Afterwards, the inoculum was aspirated and the cells were overlaid with 300 μl of 1.5% methyl-cellulose and incubated for 3 days. Then, the plates were fixed with 6% formaldehyde for 1 h followed by a 1 h staining with 300 μl crystal violet/well. After 2-3 days, the plaques were counted manually under an inverted light microscope.

### In Vivo experiments (hamster)

A total of 56 Golden Syrian hamsters, 36 male and 20 female, weighing between 80g and 130g were used in the study. Animals were weighed prior to the start of the study and randomly distributed in the different cohorts. Each of the 5 prophylactic cohorts contained 4 male and 4 female hamsters. Each of the four therapeutic cohort contained 4 male hamsters. The animals were monitored twice daily at least 6 hours apart during the study period. Body weights were measured once daily during the study period. Antibodies were diluted in PBS and dosed at the indicated concentrations in a constant volume of 500μl through intraperitoneal (IP) injection either one day before challenge with virus (“prophylaxis”) or one day after challenging with virus (“therapy”). Animals were challenged at day 0 with SARS-CoV-2 by administration of 0.05mL of a 1:10 dilution of SARS-CoV2 (OWS stock, CAT#-NR-53780), into each nostril.

### ADCC/ADCP/CDC

As target cells HEK293T cells expressing SARS-CoV-2 spike protein were used. As control cells non-transfected HEK293T and Raji B cells were used. As Effector cells human NK cells freshly isolated from human PBMC obtained under ethics permission BASEC-2016-01260.

### Comparator antibodies

Sequence information of the variable region of antibodies REGN10933 and REGN10987 was obtained from Hansen et al. Science 2020. Antibodies were expressed in HEK293 cells after gene synthesis of the variable regions and linkage to the same constant region derived from human IgG1 (Trastuzumab) as COVAB. Negative control/isotype control antibody (mAb24C3) is a human-derived antibody specific for tetanus toxoid.

## Acknowledgments

The authors want to thank our blood donors and all those who volunteered to give blood but could not be enrolled. Special thanks also to Dr. Eva Brombacher from the cantonal ethics commission for her helpful hand with the ethics approval and to Dr. Roman Diener for having accepted to be our study physician. We also thank BIOQUAL, Rockville, USA, for performing the in vivo efficacy experiment.

